# Corticostriatal dynamics underlying components of binge-like eating in mice

**DOI:** 10.1101/2021.11.04.467011

**Authors:** Britny A. Hildebrandt, Hayley Fisher, Zoe LaPalombara, Michael E. Young, Susanne E. Ahmari

## Abstract

Binge eating (BE) is a maladaptive repetitive feeding behavior present across nearly all eating disorder diagnoses. Despite the substantial negative impact of BE on psychological and physiological health, its underlying neural mechanisms are largely unknown. Other repetitive behavior disorders (e.g., obsessive compulsive disorder) show dysfunction within corticostriatal circuitry. Additionally, previous pre-clinical and clinical work has highlighted an imbalance between goal-directed and habitual responding in BE. The aim of the current study was to longitudinally examine *in vivo* neural activity within corticostriatal regions associated with habitual behavior– the infralimbic cortex (IL) and dorsolateral striatum (DLS)– in a robust pre-clinical model for BE. Female C57BL/6 mice (N=32) were randomized to receive: 1) intermittent (daily, 2-hour) binge-like access to palatable food (BE mice), or 2) continuous, non-intermittent (24-hour) access to palatable food (non-BE mice). *In vivo* calcium imaging was performed via fiber photometry at baseline and after chronic (4 weeks) engagement in the model for BE. Feeding behaviors (feeding bout onset/offset) during the recordings were captured using lickometers which generated TTL outputs for precise alignment of behavior to neural data. IL showed no specific changes in neural activity related to BE. However, BE animals showed decreased DLS activity at feeding onset and offset at the chronic timepoint when compared to baseline. Additionally, BE mice had significantly lower DLS activity at feeding onset and offset at the chronic timepoint compared to non-BE mice. These results point to a role for DLS hypofunction in chronic BE, highlighting a potential target for future treatment intervention.

**Significance Statement:** Binge eating is a chronic and repetitive eating behavior that is associated with poor physiological and psychosocial outcomes. Despite the negative impact of binge eating, little is known about the neurobiological mechanisms contributing to the chronic course and persistence of the behavior. To investigate potential neural mechanisms underlying binge eating, we are using approaches developed to monitor neural activity in rodents. This study is the first to identify longitudinal changes in neural activity within regions of the prefrontal cortex and dorsal striatum during binge-like eating behavior in mice. Findings from this work could inform targeted biological treatments for binge eating.

## Introduction

Binge eating (BE) is a chronic and repetitive eating disorder behavior that is present across nearly all eating disorder diagnoses (i.e., anorexia nervosa binge-purge subtype, bulimia nervosa, binge eating disorder, American Psychiatric Association, 2013) and is also widely prevalent in the general population. BE is strongly associated with elevated rates of obesity (Spitzer et al., 1993; Stice et al., 1999; Stice et al., 2002), poor psychosocial outcomes (e.g., suicidal ideation, Telch and Stice, 1998; Conti et al., 2017), and significant medical consequences (e.g., type II diabetes, Herpertz et al., 1998). Despite the substantial negative impact of BE, current treatments for BE and binge-related eating disorders are often inadequate, rates of relapse and persistence of eating disorder symptoms are high (Keel et al., 2005; Olmsted et al., 2015), and there has been little success with pharmacological treatments developed specifically for BE (Vocks et al., 2010; Goracci et al., 2015). Together, this underscores a critical need for research aimed at identifying the neural mechanisms contributing to BE, which could lead to the development of more effective biologically informed treatments.

Aberrant activity within corticostriatal circuitry has been found in psychiatric conditions associated with maladaptive repetitive behaviors such as obsessive compulsive disorder and substance use disorders (Graybiel and Rauch, 2000; Kalivas, 2008; Harrison et al., 2009; George and Koob, 2010; Burguiere et al., 2015). Models for these disorders have highlighted dysfunction in the balance between goal-directed and habitual behavior, suggesting increased reliance on circuitry associated with habit after more chronic durations of illness (Gillan et al., 2011; de Wit et al., 2012; Hogarth et al., 2013). However, despite evidence for corticostriatal dysfunction underlying other psychiatric conditions with prominent repetitive behaviors, there is little work directly investigating corticostriatal circuit activity in BE.

Clinical BE research has suggested that a transition to compulsive/habitual action may be present in BE related disorders (Pearson et al., 2015; Moore et al., 2016), and that aberrant function within corticostriatal circuitry may underlie BE behavior (Kessler et al., 2016). However, to date, most clinical studies attempting to investigate habit/goal-directed action and the associated neural underpinnings of BE have largely targeted obese, non-BE populations (Horstmann et al., 2015; Janssen et al., 2017). Pre-clinical work has found that male rats with a history of chronic BE show increased habitual responding during devaluation and increased activity markers (measured using c-Fos) in corticostriatal regions associated with habit – the infralimbic cortex (IL) and dorsolateral striatum (DLS) (Furlong et al., 2014). Together, these findings suggest that chronic binge-like eating may lead to habitual behavior and corresponding aberrant activity within corticostriatal circuitry.

While findings from previous work provide evidence that a history of BE leads to dysregulation of corticostriatal circuits related to habitual performance, to date, there have been no *in vivo* examinations of activity within brain regions associated with habitual behavior during binge-like eating episodes. The aim of this study was to fill this gap by identifying corticostriatal dynamics associated with food intake during BE using *in vivo* fiber photometry to capture neural activity in the IL and DLS of BE and non-BE mice. We hypothesized that after a chronic duration of binge-like eating, BE mice would show changes in neural activity within regions associated with habitual behavior (IL, DLS) during specific feeding behaviors (feeding onset, feeding offset) compared to non-BE mice, suggesting aberrant corticostriatal function after chronic BE.

## Methods and Materials

### Animals

Thirty-two adult female C57BL/6 mice (from Jackson Laboratories or in house breeding) were used for all experiments. Mice were group housed with 3-5 mice per cage and had *ad libitum* access to food and water for entirety of study. Animals were maintained on a 12/12-hour light-dark cycle (lights off at 7:00 AM; on at 7:00PM). All experiments were approved by the Institutional Animal Use and Care Committee at the University of Pittsburgh in compliance with National Institutes of Health guidelines for the care and use of laboratory animals.

### Stereotaxic Surgeries

Animals were allowed to acclimate to the light-dark cycle for approximately 10 days prior to surgery. On day of surgery, mice were anesthetized using 5% isoflurane combined with oxygen for duration of surgery. Burr holes were drilled unilaterally in left hemisphere over target regions for virus injection and fiberoptic implant. A unilateral injection of 350 nL AAV1-Synapsin.NES-jRCaMP1b.WPRE.SV40 (titer 2.0×10^12^; Addgene) and 500 nL AAV9-Synapsin-GCaMP6m-WPRE-SV40 (titer 2.7×10^12^; Addgene) were injected into IL (AP: 2.28, ML: 0.3, DV: −2.15) and DLS (AP: 0.9, ML: 2.6, DV: −2.3), respectively. Following viral injection, optical fibers (NA = .37, Neurophotometrics) were implanted into IL and DLS using the same coordinates noted above. After completion of surgery, mice were monitored until fully recovered from anesthesia, and then remained in home cage for 3 weeks to allow for recovery and viral incubation.

### Binge Eating Paradigm

The BE paradigm is based on previous work in male rats (Furlong et al., 2014), and was adapted and tested in female mice in our laboratory. We chose to focus our work on female mice given the well-established sex difference of greater incidence of BE in females vs. males in both humans (Klump et al., 2017) and rodents (Klump et al., 2013; Culbert et al., 2018). For the current study, mice were randomized to one of two feeding groups: 1) **BE**, in which animals received binge-like intermittent access to PF daily for 2-hours; or 2) **non-BE**, in which animals received continuous (24-hour), non-intermittent access to PF. The BE condition selected for this paradigm recapitulates key clinical diagnostic characteristics including intermittent consumption of a large amount of food which is consumed in a short period of time (American Psychiatric Association, 2013). While not a diagnostic requirement, food consumed during BE is typically palatable in nature–i.e., high in sweetness and fat, but low in nutritional value (Drewnowski et al., 1987; Kales, 1990; Yanovski et al., 1992; Gendall et al., 1999). In line with this, the PF used in the paradigm was sweetened condensed milk, diluted in a 3:1 ratio with water (Furlong et al., 2014). All mice had *ad libitum* access to standard chow throughout the paradigm and were never placed on food restriction.

Mice completed four weeks of daily PF feeding tests. Prior to each feeding test, mice were removed from group-housed conditions and placed in individual feeding test cages for duration of each 2-hour feeding test. PF (delivered via 50mL conical tube with sipper attachment) and standard chow were weighed at the beginning and end of the 2-hour feeding test, and body weight was measured daily. At completion of each feeding test, mice were removed from feeding test cages and returned to group housed home cage. Animals in the non-BE group had a new bottle of PF placed in the home cage daily for continuous access to PF. This bottle was weighed daily prior to feeding test.

### Fiber Photometry Recordings and Processing

All *in vivo* fiber photometry recordings took place in operant chambers (Med Associates) equipped with contact lickometers mounted to chamber wall. Contact lickometers (Med Associates) registering TTL outputs for each individual lick were used to identify feeding onset (first lick of a feeding bout) and feeding offset (last lick of a feeding bout). Bouts were defined as ≥ 2 licks with breaks in between licking of no greater than 3 seconds. Prior to the start of the study, all animals were habituated over multiple days to the operant chamber and lickometer (containing only water). Animals were also habituated for at least two sessions to scruffing and cable attachment. There were two photometry recordings during the study– one at a *baseline* timepoint (prior to initiation of BE paradigm) and one at a *chronic* timepoint (after four weeks of engagement in BE paradigm). Animals were water restricted (and PF restricted for non-BE group) overnight prior to recording days, while maintaining *ad libitum* access to standard chow. The first recording at the baseline timepoint occurred prior to initiation of BE paradigm, and animals were therefore naïve to PF. This design permits monitoring activity from first exposure/onset through chronic duration of behavior. The baseline recording was 40 minutes in duration to account for neophobia to PF and ensure enough feeding bouts for meaningful interpretation of neural data. The second recording was 20 minutes and took place at chronic timepoint after completion of BE paradigm (i.e., four weeks).

Photometry recordings used a 3-color, 2-region system (Neurophotometrics). Animals were connected using a 4x branching cable (Doric) to allow for simultaneous recording from 2 regions in each animal. The system pulsed at 40Hz and interleaved three LED channels (415nm, 470nm, 560nm) during recordings, resulting in detection of 1) isosbestic signal, 2) GCaMP6m activity (DLS), and 3) jRCaMP1b activity (IL). Bonsai was used to interface photometry system with Med Associates TTL data and provide synchronization with photometry signal in real time. After recordings, traces were separated to examine activity within each channel independently. All photometry data was processed and analyzed using custom MATLAB scripts. Processing began by removing motion artifacts from 470nm (GCaMP6m) and 560nm (jRCaMP1b) channels using the 415nm channel (isosbestic) trace as a reference point. Next, traces were corrected for exponential decay and run through a low-pass filter (order 6, frequency 3). A moving minimum using a sliding minimum of 120 seconds was applied to remove large fluctuations from the traces. Finally, traces were z-scored for normalization. These traces were aligned with TTL outputs from the lickometers for analysis of neural activity at onset and offset of feeding bouts. For each trace, data were aligned to behavior of interest at timepoint “0” (feeding onset, feeding offset), and trace analysis window included three seconds prior to the event and one second after the event.

### Histology

After completion of BE paradigm and photometry recordings, all mice were transcardially perfused using 4% paraformaldehyde (PFA). Brains were removed and remained in PFA for an additional 24 hours, cut into 35-micron sections, mounted onto slides, and cover slipped with DAPI mounting media. All sides were scanned using an Olympus slide scanning microscope. Location of fiber implant was identified by examining damage tracks in tissue, and placement of fiber and spread of virus were used to include or exclude animals from analysis (Figure 1A; Extended Data Figure 1-1).

**Figure 1.**
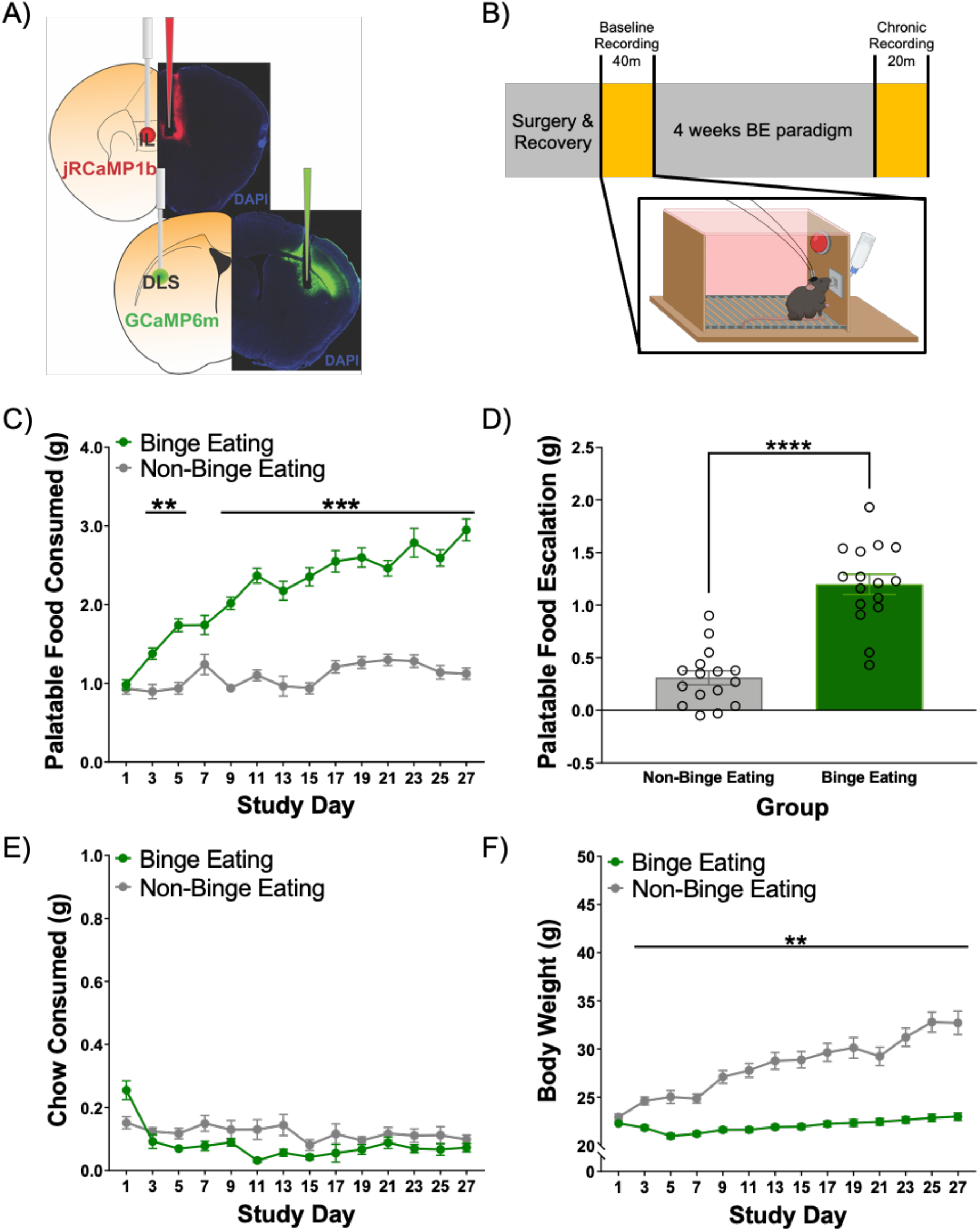
Binge eating paradigm behavioral data. A) Mice were injected unilaterally in left hemisphere with AAV1-Synapsin.NED-jRCaMP1b.WPRE.SV40 into IL and AAV9-Synapsin-GCaMP6m-WPRE-SV40 into DLS. Optical fibers were implanted into IL and DLS after viral injection. B) Study timeline. Inset: schematic of *in vivo* recording setup. For behavior data, palatable food (PF), chow, and body weight (BW) data are graphed showing every other day. N = 16 BE, 16 Non-BE. C) BE animals consumed significantly more PF compared to non-BE animals during feeding tests. D) Only BE animals escalated their PF intake across the study period. E) There were no specific days identified where chow intake was different between BE and non-BE mice. F) Body weight significantly increased in non-BE group across the study period. **p < 0.01, ***p < .001, ****p < .0001.

### Statistical Analyses

For data from BE paradigm, repeated measures ANOVAs (or mixed models in case of missing data) with Bonferroni post-hoc analyses or individual samples t-tests were used. Results are presented as mean ± SEM. A p-value of ≤ 0.05 was set to determine statistical significance. Statistical analyses were run using GraphPad Prism 9.

For fiber photometry data, multilevel spline regressions were run on normalized photometry data during feeding bouts (group analysis of IL and DLS data) or the residuals of the best fitting model (exploratory variance analysis of DLS data). The best fitting models were chosen via model comparisons that employed the Bayesian information criterion (BIC), which considers the degree of flexibility (i.e., number of change points) of the model and penalizes more flexible models (Kuha, 2004). All models were a full factorial analysis, and all categorical predictors were effect coded. For all analyses, the residual distributions were examined for normality and homoscedasticity. All group analysis models met assumptions, but the two variance analysis models were heteroscedastic. To satisfy the assumption of homoscedasticity, the residuals of the best fitting models were square root transformed prior to analysis. Predictors included the between-subjects categorical variable Group (BE, non-BE), and the within-subjects categorical variable Stage (Baseline, Chronic) and continuous variable Time (−3 seconds prior to event of interest to 1 second after event of interest). The random effects structure included Intercept (subject) and two slopes (Stage and Time). Planned comparisons were run for significant interactions including Time, Stage, and Group using Type III Sums of Squares. All photometry data were analyzed using *R* (R 4.0.3). The highest order significant interactions (*p* < 0.05) including the variables Stage and Group were followed by planned comparisons using *R*’s emmeans package. Group × Stage × Time interactions were followed by planned comparisons at 500 ms intervals using emmeans. Results are presented as the model fit mean ± SEM. Raw values alongside model fits are presented in Extended Data Tables 2-1 and 3-1.

## Results

### Binge-like eating leads to escalated PF intake

All animals completed four weeks of the BE paradigm (Figure 1B). On feeding test days, total intake of PF and chow during each feeding test was measured. Body weight was also measured daily. For PF intake, there was a significant Time x Group interaction (F(8.32, 237.4) = 31.60, *p* < .0001) such that BE animals consumed significantly more PF compared to non-BE mice during feeding tests starting on Day 2 (Figure 1C). This pattern persisted for the duration of the study (all *ps* < .001; except for Day 2, 3: *p* < 01; Day 7: *p* = .21). BE animals also escalated their PF intake (average PF intake week 4 – average PF intake week 1) during feeding tests significantly more than non-BE mice across the four week BE paradigm (Figure 1D; t(30) = 7.63, *p* < .0001). There was a significant Time x Group interaction for chow intake during feeding tests (Figure 1E, F(26, 764) = 2.67, *p* < .0001); however, post-hoc analysis identified no significant differences across days between BE and non-BE mice. There was also a significant Time x Group interaction for body weight (Figure 1F, F(26, 780) = 50.65, *p* < .0001), with non-BE mice weighing significantly more than BE animals; this difference emerged on study Day 2 and persisted across study (all *p’s* < .01). Figure 1 shows data plotted every other day across study; Extended Figure 1-2 shows daily data.

### Binge eating paradigm does not lead to significant differences in IL activity measured using fiber photometry

Comparing Bayesian Information Criteria (BICs), the best fitting model for IL feeding onset included five change points. There was no significant Group × Stage (χ^2^(1) = 0.05, *p* = 0.83) or Group × Stage × Time (χ^2^(5) = 1.69, *p* = 0.89) interaction. There were no differences in activity between or within BE and non-BE groups (Figures 2A, 2B). Other lower order effects are shown in Extended Data Table 2-1. Comparing BICs, the best fitting model for IL feeding offset included six change points. There was a significant Group × Stage × Time interaction (χ^2^(5) = 26.24, *p* < 0.01). The interaction revealed that the non-BE group had reduced activity compared to baseline at −1.0 seconds (*z* = 2.42, *p* = 0.02) prior to feeding offset (Figure 2C). There were no differences within the BE group (all *p* > 0.10). At baseline, BE animals had lower activity than the non-BE group at −1.5 (*z* = −2.27, *p* = 0.03) and −1.0 (*z* = −2.57, *p* = 0.02) seconds prior to feeding offset (Figure 2D). At the chronic timepoint, BE animals had lower activity at −2.5 (*z* = −2.27, *p* = 0.03), −2.0 (*z* = −2.85, *p* < 0.01), and −1.5 (*z* = −2.75, *p* = 0.01) seconds compared to the non-BE group (Figure 2D). However, these results at the chronic timepoint are inconclusive given significant differences in baseline activity between groups (Figure 2D, left panel). Other lower order effects are shown in Extended Data Table 2-1.

**Figure 2.**
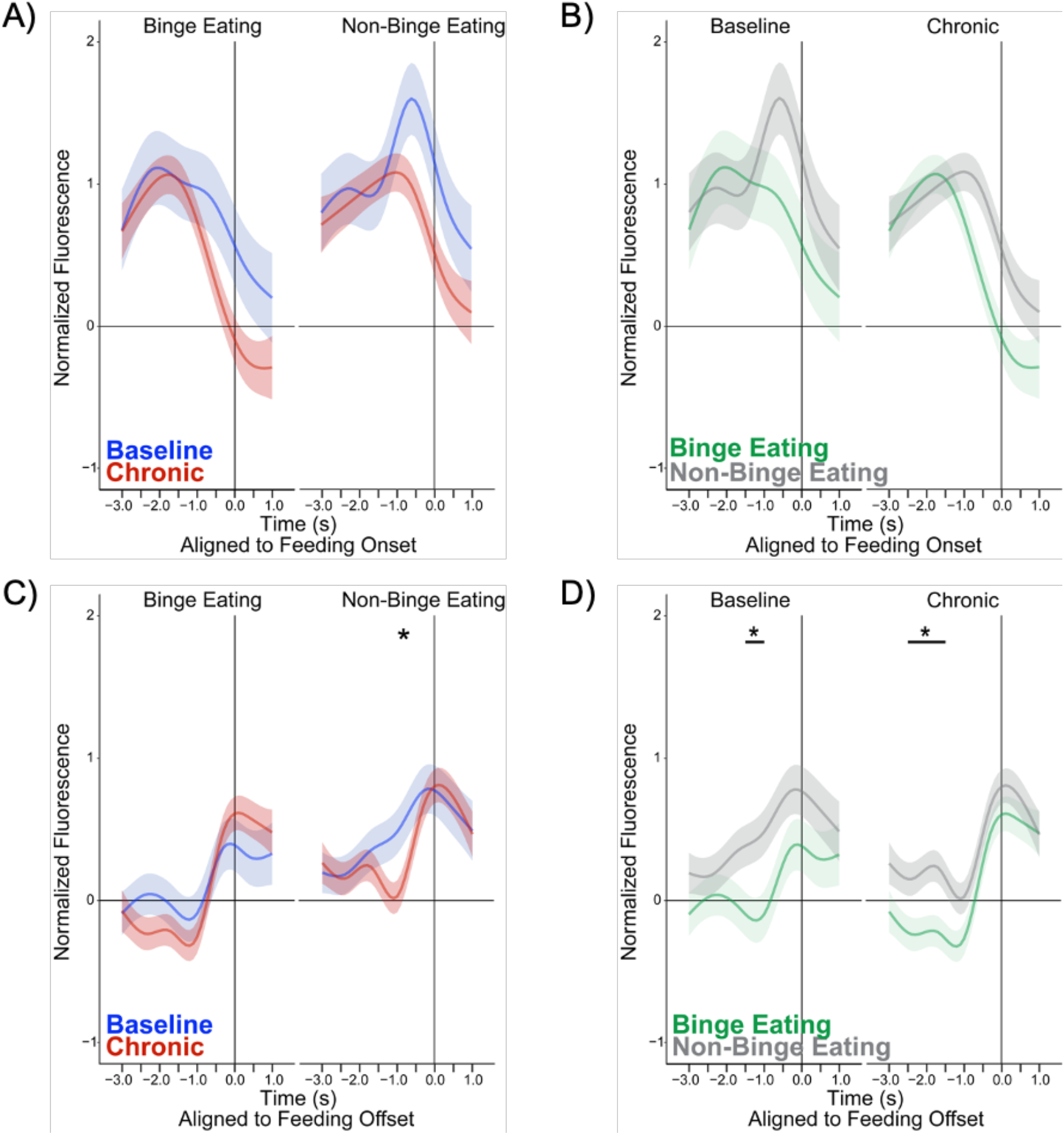
In vivo fiber photometry in IL at feeding onset and offset. All photometry data are aligned to feeding onset (A, B) or feeding offset (C, D) at Time = 0. Figures A and C represent within group analyses. Figures B and D represent between group analyses. The same data are plotted in two different ways to visualize effects for inspection by the reader. A) There were no differences within BE or non-BE groups in IL activity associated with feeding onset. B) There were no differences between BE and non-BE mice in feeding onset-associated IL activity. C) Non-BE animals had significantly lower activity at the chronic timepoint −1.0 seconds prior to feeding offset vs. baseline. D) At baseline, BE animals had significantly lower activity vs. non-BE animals −1.5 and 1.0 seconds before feeding offset. At the chronic timepoint, BE animals had significantly lower activity −2.5, −2.0, and −1.5 seconds before feeding offset vs. non-BE animals. *p < 0.05.

### Binge-like eating leads to decreased DLS activity at Feeding Onset

Comparing BICs, the best fitting model for DLS feeding onset included five change points. There was a significant Group × Stage × Time interaction (χ^2^(5) = 41.17, *p* < 0.01). The BE group had reduced DLS activity at the chronic timepoint compared to baseline at −2.5 (*z* = 2.13, *p* = 0.03) and −2.0 (*z* = 2.16, *p* = 0.03) seconds prior to feeding onset (Figure 3A). There were no significant differences between baseline and chronic timepoints in non-BE animals (Figure 3A). Between group analysis showed that, compared to the non-BE group, BE animals had reduced activity from −2.5 to −1.0 seconds (*z*_s_ = −4.46 to −2.72, all *p* < 0.01) prior to feeding bout onset at the chronic timepoint (Figure 3B). Other lower order effects can be found in Extended Data Table 3-1. Exploratory analyses examining variance across groups found a significant Group × Stage interaction (χ^2^(1) = 4.32, *p* = 0.04). The BE group had reduced variability in their neural activity related to feeding onset at the chronic timepoint compared to baseline (*z* = 2.29, *p* = 0.02) (Figure 3C), while there were no changes in variance in the non-BE group (*z* = −1.27, *p* = 0.21) (Figure 3C). The interaction also showed that at the chronic timepoint, the BE group had lower variability than the non-BE group (*z* = −2.34, *p* = 0.02); this difference was not observed during the baseline timepoint (*z* = 1.17, *p* = 0.24) (Figure 3C).

**Figure 3.**
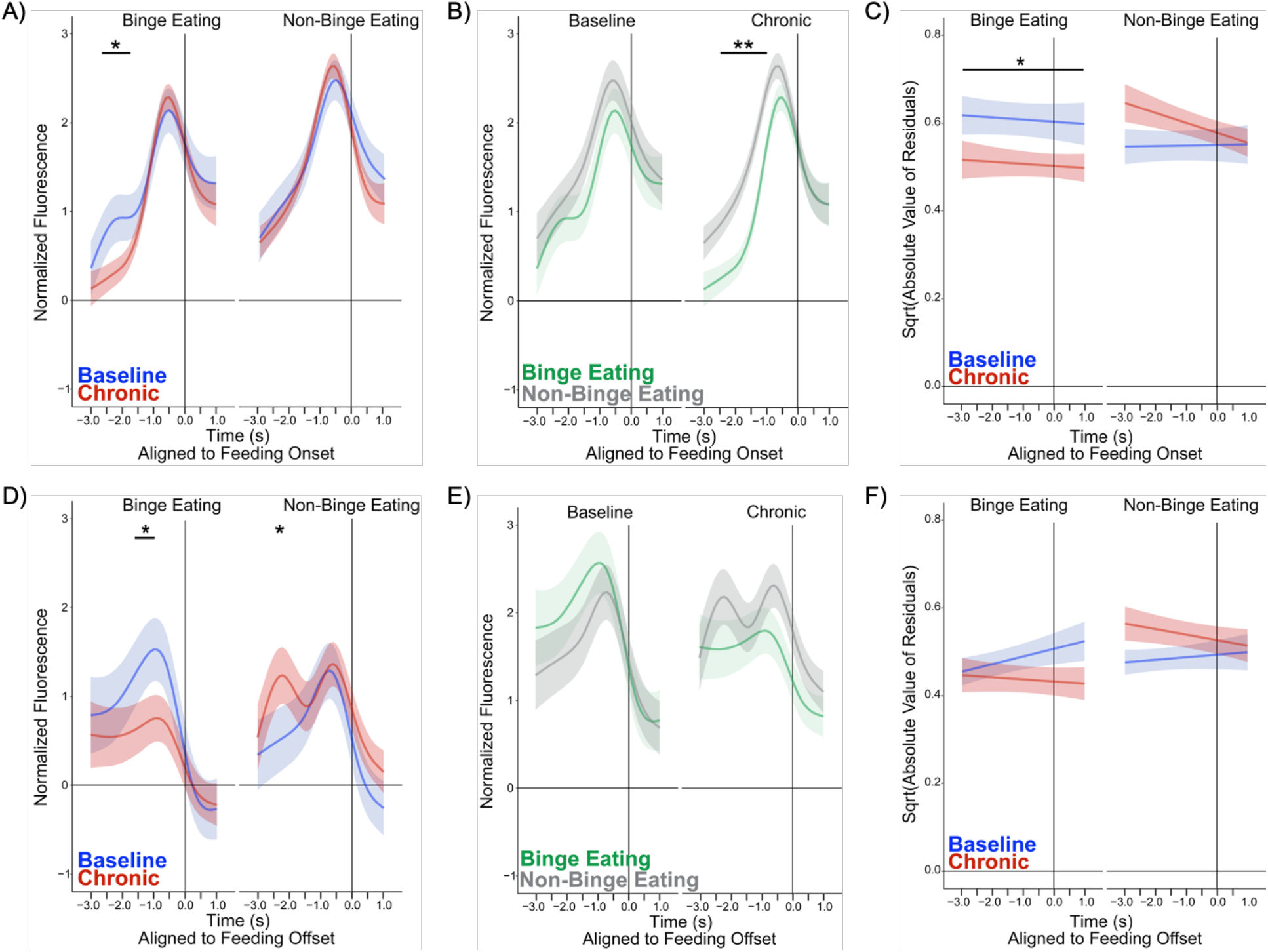
In vivo fiber photometry in DLS at feeding onset and offset. All data are aligned to feeding onset (A, B, C) or feeding offset (D, E, F) at Time = 0. Figures A and D represent within group analysis. Figures B and E represent between group analysis. The same data are plotted in two different ways to visualize effects for inspection by the reader. A) BE mice had significantly lower activity at chronic timepoint vs. baseline at −2.5 and −2.0 seconds prior to feeding onset. B) BE animals had significantly lower activity vs. non-BE animals at the chronic timepoint −2.5 to −1.0 seconds prior to feeding onset. C) Variance in activity associated with feeding onset significantly decreased from the acute to chronic timepoint in BE, but not in non-BE mice. Variance at chronic timepoint was lower in BE vs. non-BE mice. Group × Stage interaction (c^2^(1) = 4.25, *p* = 0.04). D) BE mice had significantly lower activity at chronic vs. baseline timepoint at −1.5 and −1.0 seconds prior to feeding offset. Non-BE mice had significantly higher activity at chronic timepoint vs. baseline at −2.5 seconds. E) Trend toward significant difference at chronic timepoint with lower activity at −0.5 and 0.0 seconds before feeding offset in BE vs. non-BE mice. F) No differences in variance related to feeding offset in BE and non-BE mice. *p < 0.05, **p < 0.01.

### Trend towards decreased DLS activity at Feeding Offset after binge-like eating

Comparing BICs, the best fitting model for DLS feeding offset included five change points. There was a significant Group × Stage × Time interaction (χ^2^(5) = 30.51, *p* < 0.01). The BE group had reduced activity in DLS at the chronic timepoint compared to baseline at −1.5 (*z* = 1.97, *p* = 0.048) and −1.0 (*z* = 2.33, *p* = 0.02) seconds prior to feeding offset (Figure 3D). The non-BE group had increased activity at the chronic timepoint compared to baseline at −2.5 (*z* = −2.13, *p* = 0.03) seconds (Figure 3D). Compared to the non-BE group, BE animals trended towards having reduced activity at −0.5 (*z* = −1.80, *p* = 0.07) and 0 (*z* = −1.76, *p* = 0.08) seconds prior to feeding bout offset at the chronic, but not baseline, timepoint (Figure 3E). Other lower order effects are shown in Extended Data Table 3-1. Exploratory analyses examining variance in DLS activity associated with feeding offset found a significant Group × Stage interaction (χ^2^(1) = 5.64, *p* = 0.02). The interaction revealed no within group differences between timepoints, but showed that at the chronic timepoint, the BE group had lower variability than the non-BE group (*z* = −2.39, *p* = 0.02); this was not observed at the baseline timepoint (*z* = 0.08, *p* = 0.94) (Figure 3F).

## Discussion

This study used *in vivo* fiber photometry to investigate longitudinal changes in neural activity within corticostriatal circuitry in BE and non-BE mice. We examined activity during naturalistic binge-like eating episodes aligned to specific feeding behaviors (feeding onset, feeding offset) that can also be quantitatively measured in humans. While IL activity showed no specific changes after undergoing the BE paradigm (Figure 2; observed differences in BE group are present at both baseline and chronic timepoints), we found differential activity patterns in the DLS between BE and non-BE mice after a chronic duration of PF intake (Figure 3). Additionally, findings highlighted that a history of binge-like eating leads to a stronger impact on feeding onset rather than offset in DLS (comparison of Figs. 3B and E). Together, this suggests that reduced recruitment of DLS, particularly during feeding onset, is specific to animals with a history of binge-like eating (i.e., intermittent) and not general (i.e., continuous) PF intake. These results point to a neural mechanism in the DLS underlying chronic BE, highlighting a potential target for future treatment intervention.

Previous work has suggested that BE leads to an imbalance in goal-directed and habitual behavior. Clinical research of BE populations has found that when viewing images of PF, individuals with BE show increased recruitment of brain regions involved in goal-directed behavior (e.g., ventromedial prefrontal cortex, Neveu et al., 2018). However, on a decision making task unrelated to food, individuals with BE disorder engaged in significantly more habitual responding (Voon et al., 2015). Together, these findings suggest that feeding and food related tasks may differently recruit goal-directed/habitual circuitry compared to tasks assessing habitual decision-making (unrelated to feeding and food) in individuals with BE. In pre-clinical studies, baseline deficits in goal-directed responding in rats were associated with higher levels of subsequent binge-eating upon exposure to PF (LeMon et al., 2019), suggesting that impaired goal-directed responding may be a risk factor for BE development. Other work has shown that rats with a history of BE engaged in more lever pressing during outcome devaluation testing, suggesting BE led to more habitual responding compared to continuous access or control rats (Furlong et al., 2014). This pattern of habitual responding was associated with increased outcome-devaluation-test-related-c-Fos activity in areas associated with habit (i.e., DLS, IL, Furlong et al., 2014). This contrasts with our findings of decreased DLS activity after chronic BE at feeding onset and offset; thus, our findings suggest that in our paradigm, BE-related feeding behaviors are not driven by habit. However, the current study examined neural activity in real time rather than using an indirect marker of neural activity like c-Fos, which may explain the differences in results. Nonetheless, future work should investigate engagement of corticostriatal regions associated with goal-directed behavior (e.g., prelimbic cortex, dorsomedial striatum) during binge-like eating to help disentangle the relationship between goal-directed and habitual behavior in control of pathological BE feeding behavior.

Feeding bouts involve a repetitive sequence of behaviors including feeding onset, consumption, and feeding offset, and previous work has shown that DLS plays a key role in initiating and completing sequences. Training on a 2-step lever pressing task showed that DLS lesions impaired acquisition of correct sequence completion in mice (Yin, 2010), and work using a viral strategy to specifically suppress activity of D1+ spiny projection neurons in DLS found impairments in sequence accuracy (Rothwell et al., 2015). We measured DLS activity during distinct phases of the feeding sequence (feeding onset, feeding offset). At both feeding onset and offset, BE animals showed a reduction in DLS activity from their baseline to the chronic timepoint. At the chronic timepoint, BE animals also had lower activity compared to non-BE animals at feeding onset (Figure 3B). At feeding offset, BE mice showed a blunting of activity from baseline to the chronic timepoint (Figure 3D) and trended toward being significantly lower than non-BE mice at the chronic timepoint (Figure 3E). Our findings showing decreased/blunted DLS activity in BE mice may contribute to difficulty in executing proper feeding sequences in BE mice.

Reduced activity in DLS of BE mice may result from decreased activity in upstream regions that project to the DLS and contribute to sequence performance, such as secondary motor cortex (M2), a primary input to DLS (Rothwell et al., 2015; Lee et al., 2019). Pharmacological inactivation of M2 in rats with a history of BE increased PF consumption (Corwin et al., 2016), suggesting that lower M2 activity may contribute to impairments in feeding offset/feeding sequence completion. M2→DLS projections are critical for correct execution and termination of behavioral sequences (Rothwell et al., 2015). Therefore, the blunted activity found at feeding onset in DLS of BE mice may represent less input from M2→DLS projections, thus promoting ongoing repetitive action. While these data underscore the importance of DLS and associated inputs in generating sequenced behavior, future work should use alternative sequence testing paradigms in chronic BE and non-BE mice to determine if these findings are unique to BE or generalize to other sequenced behaviors. Additionally, manipulation of M2→DLS projections during distinct phases of the feeding sequence will help directly dissect the role of these projections in binge-like eating.

In conclusion, here we capitalized on pre-clinical approaches to longitudinally examine *in vivo* neural activity associated with feeding behaviors during binge-like eating. Our results demonstrate decreased calcium activity in the DLS after chronic BE, pointing to a potential target for development of biologically informed treatment interventions. Future work should further investigate corticostriatal contributions to BE behavior and the interplay between goal-directed and habitual behavior in this context.

## Acknowledgements

This work was supported by the BRAIN Initiative and the National Institutes of Mental Health (F32MH118687, BH; T32MH016804).

---

**Extended Data Table 2-1.** Lower level photometry effects for infralimbic cortex. Categorical predictors are effect coded. Bold indicates *p* < 0.05.

| *IL Activity at Feeding Onset* | $\chi^2$ | df | *p* |
| --- | --- | --- | --- |
| Intercept | 24.35 | 1 | **< 0.01** |
| Time (df = 5) | 321.94 | 5 | **< 0.01** |
| Stage | 0.07 | 1 | 0.79 |
| Group | 0.09 | 1 | 0.77 |
| Time*Stage | 1114.19 | 5 | **< 0.01** |
| Time*Group | 68.90 | 5 | **< 0.01** |
| Stage*Group | 0.05 | 1 | 0.83 |
| Time*Stage*Group | 1.69 | 5 | 0.89 |
| *IL Activity at Feeding Offset* | $\chi^2$ | df | *p* |
| Intercept | 0.96 | 1 | 0.33 |
| Time (df = 6) | 255.25 | 6 | **< 0.01** |
| Stage | 0.09 | 1 | 0.77 |
| Group | 4.86 | 1 | **0.03** |
| Time*Stage | 74.77 | 6 | **< 0.01** |
| Time*Group | 21.32 | 6 | **< 0.01** |
| Stage*Group | 0.03 | 1 | 0.87 |
| Time*Stage*Group | 26.24 | 6 | **< 0.01** |

**Extended Data Table 3-1.** Lower level photometry effects for dorsolateral striatum. Categorical predictors are effect coded. Bold indicates *p* < 0.05.

| *DLS Activity at Feeding Onset* | $\chi^2$ | df | *p* |
| --- | --- | --- | --- |
| Intercept | 9.59 | 1 | **< 0.01** |
| Time (df = 5) | 2436.71 | 5 | **< 0.01** |
| Stage | 0.59 | 1 | 0.44 |
| Group | 2.08 | 1 | 0.15 |
| Time*Stage | 82.40 | 5 | **< 0.01** |
| Time*Group | 57.55 | 5 | **< 0.01** |
| Stage*Group | 0.24 | 1 | 0.62 |
| Time*Stage*Group | 41.17 | 5 | **< 0.01** |
| *DLS Activity at Feeding Offset* | $\chi^2$ | df | *p* |
| Intercept | 40.89 | 1 | **< 0.01** |
| Time (df = 5) | 1180.57 | 5 | **< 0.01** |
| Stage | < 0.01 | 1 | 0.97 |
| Group | 0.40 | 1 | 0.53 |
| Time*Stage | 133.17 | 5 | **< 0.01** |
| Time*Group | 44.09 | 5 | **< 0.01** |
| Stage*Group | 1.08 | 1 | 0.30 |
| Time*Stage*Group | 30.51 | 5 | **< 0.01** |

**Extended Figure 1-1.**
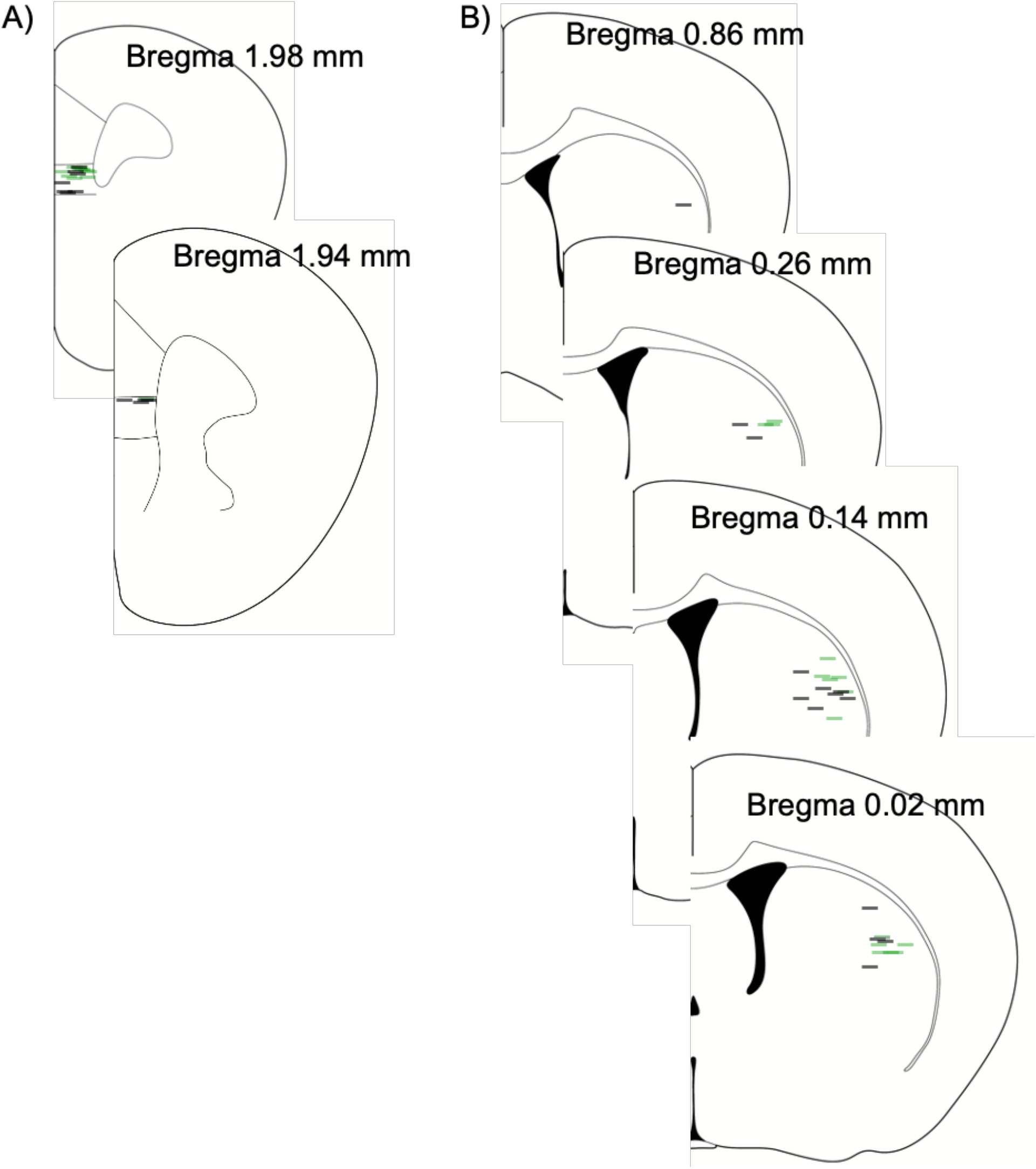
Fiber placements for in vivo fiber photometry. Histological verification of fiber placements in the A) infralimbic cortex and B) dorsolateral striatum. Each line represents one animal, and lines represent bottom of fiber. Green lines = BE, gray lines = non-BE.

**Extended Figure 1-2.**
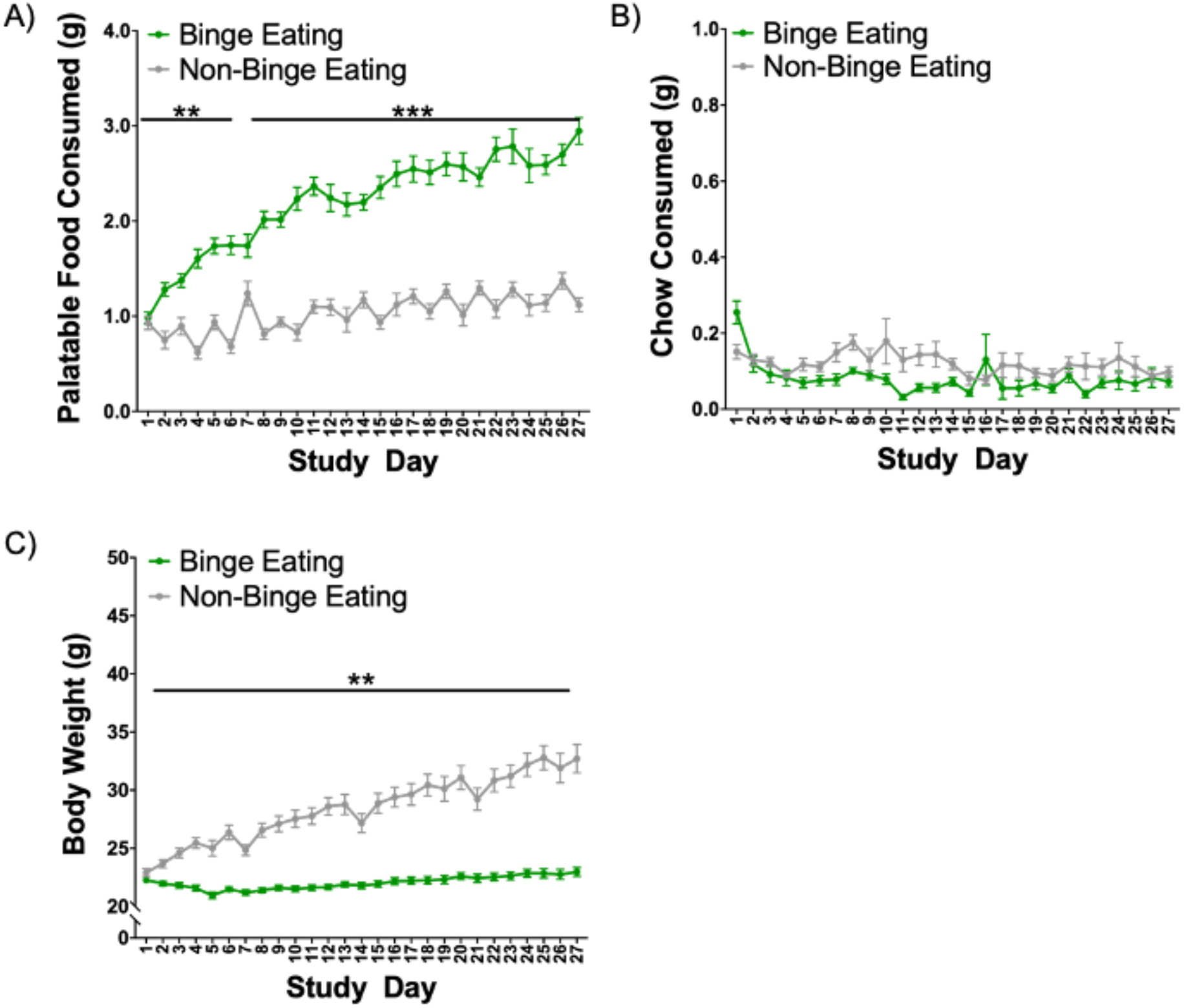
Binge eating paradigm behavioral data plotted for all study days. PF, chow, and BW data are graphed for each day of study period. N = 16 BE, 16 Non-BE. A) BE animals consumed significantly more PF compared to non-BE animals during feeding tests. B) No specific days were identified with different chow intake between BE and non-BE mice. C) Body weight significantly increased in non-BE group across the study period. **p < 0.01, ***p < .001.

